# Reduced structural connectivity in Insomnia Disorder

**DOI:** 10.1101/510784

**Authors:** Kira V. Jespersen, Angus Stevner, Henrique Fernandes, Stine D. Sørensen, Eus Van Someren, Morten Kringelbach, Peter Vuust

**Author notes:** **Corresponding author:** Kira Vibe Jespersen, Center for Music in the Brain, Noerrebrogade 44, build. 10G, 4^th^ floor, 8000 Aarhus C, Denmark.

## Abstract

Insomnia Disorder is the most prevalent sleep disorder, and it involves both sleep difficulties and daytime complaints. The neural underpinnings of Insomnia Disorder are poorly understood. Several existing neuroimaging studies focused on local measures and specific regions of interests, which makes it difficult to judge their whole-brain significance. We therefore here applied a data-driven approach to assess differences in whole-brain structural connectivity between adults with Insomnia Disorder and matched controls without sleep complaints. We used diffusion tensor imaging and probabilistic tractography to assess whole-brain structural connectivity and examined group differences using Network-Based Statistics. The results revealed a significant difference in the structural connectivity of the two groups. Participants with Insomnia Disorder showed reduced connectivity in a subnetwork that included mainly fronto-subcortical connections with the insula as a key region. By taking a whole-brain network perspective, our study enables the integration of previous inconsistent findings. Our results reveal that reduced structural connectivity of the left insula and the connections between frontal and subcortical regions are central neurobiological features of Insomnia Disorder. The importance of these areas for interoception, emotional processing, stress responses and the generation of slow wave sleep may help guide the development of neurobiology-based models of the prevalent condition of Insomnia Disorder.

---

Insomnia is the most common sleep disorder, reported in 6% to 22% of the general population^1,2^ The diagnostic criteria involve prolonged difficulties initiating or maintaining sleep, including early morning awakenings, and significant impairments in important areas of daytime functioning (e.g. occupational, social or behavioural)^3^. Unfortunately, insomnia is often a persistent condition^4,5^, and it is associated with reduced quality of life as well as deficiencies in cognitive and emotional functioning^6^. There is a high rate of comorbidity with medical and psychiatric disorders, and insomnia confers an increased risk for depression and anxiety^7–10^.

Even though the burden of insomnia is heavy for both individuals and society, the pathophysiology of the disorder is poorly understood. Different models of insomnia have been proposed^11^. The most prevailing theory of insomnia is the hyperarousal theory, which conceptualizes insomnia as the result of increased physiological and cortical arousal that interfere with the normal sleep processes^12–14^. Most neuroimaging studies have been interpreted within this framework. However, not all studies fit this model, and a more complex understanding may be needed, e.g. including awareness^15,16^ and insufficient overnight downregulation of emotional distress^17^ as key dimensions.

So far, no reliable biological marker for insomnia has been identified. Modern neuroimaging methods allow for a non-invasive mapping of structural and functional brain networks and provide a promising way to shed light on the neurobiological foundations of insomnia to achieve clinically relevant information that can help target interventions. An increasing number of studies have investigated functional connectivity alterations related to insomnia^18–27^, but as a number of recent reviews point out, the results are relatively inconsistent so far^15,16,28^. This inconsistency may stem from the variety of methods used^28^, but may also be influenced by sleep confounds^29^ and insomnia heterogeneity^30^.

Other studies have approached the neurobiological basis of insomnia by looking at structural differences between persons with Insomnia Disorder and good sleeper controls. In terms of grey matter alterations, neuroimaging studies have linked insomnia to grey matter decreases in the orbitofrontal cortex^31,32^ and hippocampus^33,34^ In addition, two studies have used diffusion tensor imaging (DTI) to evaluate differences in white matter tracts in persons with Insomnia Disorder compared to matched controls. One study found reduced fractional anisotropy in the anterior internal capsule, suggesting disturbed fronto-subcortical connectivity in patients with insomnia^35^. Similarly, Li and colleagues found reduced fractional anisotropy in the right anterior and posterior internal capsule, as well as in white matter tracts of the superior corona radiate, longitudinal fasciculus, thalamus and the corpus callosum^36^. A recent DTI study assessed topological alterations in Insomnia disorder and found changes of the regional organization of frontal and subcortical areas as well as reduced connectivity in frontal networks^37^. In summary, studies assessing structural alterations related to insomnia have generally reported reductions in frontal and subcortical areas and the white matter tracts connecting these areas.

Across functional and structural studies, a recent coordinate-based meta-analysis integrating neuroimaging findings showed limited consistency in identifying “where” in the brain findings on insomnia converge^38^. Therefore, a more integrated approach may be required to better understand the brain mechanisms of insomnia. Most studies on brain structure have assessed insomnia related differences in specific regions and white matter tracts, and the few studies that have assessed insomnia related structural connectivity, have focused mainly on graph-theoretical measures^37,39^. Existing studies on functional connectivity in insomnia have focused on seed-based methods with regions of interest defined in advance based on regionally specific hypotheses, thus disregarding potential insomnia-related reconfigurations at the whole-brain network level, for example resulting from possible axonal vulnerability^40^. The brain is an extraordinarily complex system, and modern understandings of neuropathology have evolved from an emphasis on specific brain regions to evaluating the disturbances of inter-connected neural networks^41,42^. State-of-the-art neuroimaging methods reflect this refined understanding of brain structure and function by applying data-driven connectivity approaches at the whole-brain level. This is particularly relevant to disorders such as insomnia that affects a large range of functions not plausibly explained by alterations of single brain structures. Uncovering the large-scale network alterations in insomnia could prove essential for a better understanding of this highly prevalent disorder and the widespread dysfunctions associated with it. However, this perspective has not gained sufficient attention in insomnia research.

We here employed a data-driven approach to investigate the changes in structural connectivity (SC) associated with Insomnia Disorder. SC forms the anatomical backbone of functional brain connectivity, and DTI techniques have enabled us to non-invasively assess the white matter tracts connecting brain areas and their alterations in neurological and psychiatric disorders^43^. This approach allowed us to assess whole-brain structural connectivity in adults with Insomnia Disorder compared to healthy controls with no sleep complaints. Based on the before mentioned studies, we hypothesized that Insomnia Disorder would be characterized by reduced SC among frontal and subcortical areas, and that individual differences in connectivity would correlate with insomnia severity.

## Methods

### Participants

We included 30 participants in the study; 16 persons with Insomnia Disorder and 14 matched controls without sleep complaints. Insomnia participants were recruited from the sleep clinic at the Department of Clinical Neurophysiology, Aarhus University Hospital (Denmark) or via newspaper announcements. Age- and gender-matched controls with no sleep problems were recruited through local announcements. All participants underwent a clinical interview, and inclusion criteria for the insomnia group were Insomnia Disorder according to the DSM-5 criteria. That is, difficulties initiating or maintaining sleep, including early morning awakenings, with adequate opportunity for sleep. The sleep problems had to be present for at least three nights per week for the last three months and had to be associated with significant impairments in important areas of daytime functioning, such as occupational, educational or social functioning^3^. Participants with Insomnia Disorder were screened for other sleep disorders and underwent one night of ambulatory polysomnography. They were excluded if they had more than mild symptoms of other sleep disorders, such as sleep-disordered breathing, sleep-related movement disorder or circadian rhythm disorder. Exclusion criteria for all participants were the use of psychotropic or hypnotic medications, sleep disruptive medical disorders, psychiatric disorders, alcohol or substance abuse, as well as any standard MRI incompatibility. All participants completed the Pittsburgh Sleep Quality Index (PSQI)^44^ and the Insomnia Severity Index (ISI)^45^. The characteristics of the participants are shown in Table 1. The study was approved by the Ethical Committee of the Central Denmark Region, and participants signed informed consent prior to inclusion in accordance with the Declaration of Helsinki.

**Table 1.**
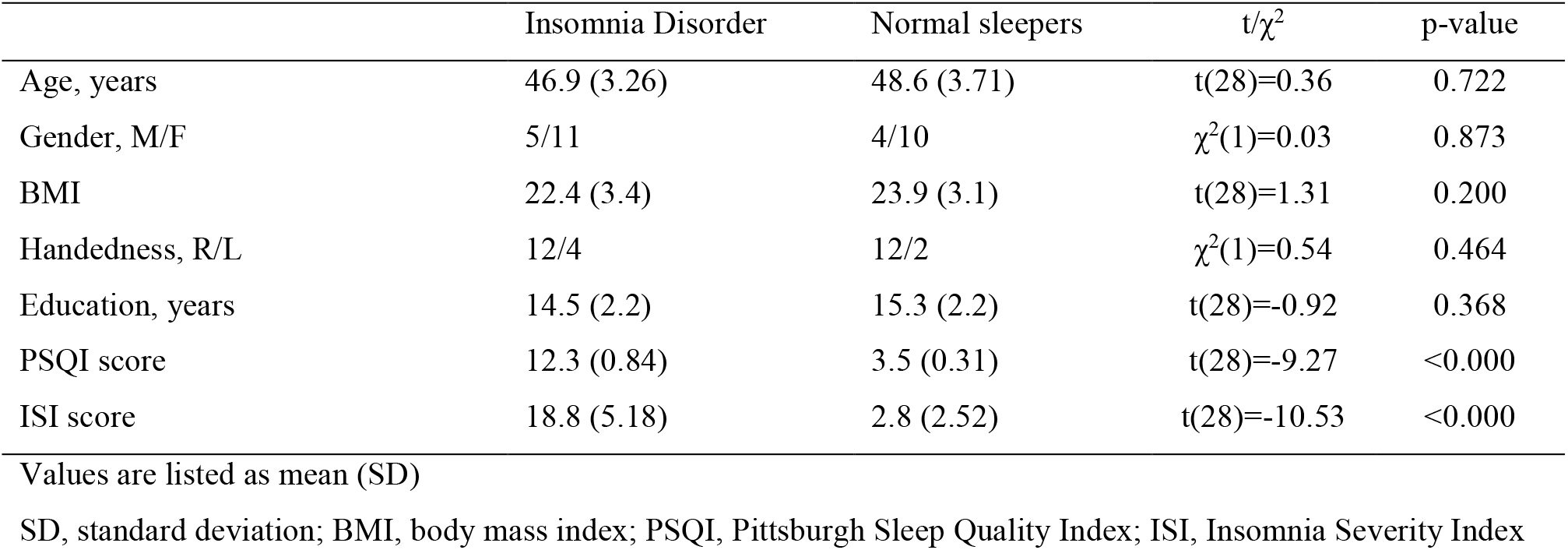
Demographic and clinical characteristics of the participants

| | Insomnia Disorder | Normal sleepers | t/$\chi^2$ | p-value |
| --- | --- | --- | --- | --- |
| Age, years | 46.9 (3.26) | 48.6 (3.71) | t(28)=0.36 | 0.722 |
| Gender, M/F | 5/11 | 4/10 | $\chi^2(1)=0.03$ | 0.873 |
| BMI | 22.4 (3.4) | 23.9 (3.1) | t(28)=1.31 | 0.200 |
| Handedness, R/L | 12/4 | 12/2 | $\chi^2(1)=0.54$ | 0.464 |
| Education, years | 14.5 (2.2) | 15.3 (2.2) | t(28)=-0.92 | 0.368 |
| PSQI score | 12.3 (0.84) | 3.5 (0.31) | t(28)=-9.27 | <0.000 |
| ISI score | 18.8 (5.18) | 2.8 (2.52) | t(28)=-10.53 | <0.000 |

Values are listed as mean (SD)
SD, standard deviation; BMI, body mass index; PSQI, Pittsburgh Sleep Quality Index; ISI, Insomnia Severity Index

### Image acquisition and analysis

All participants underwent the same imaging protocol using a Siemens Trio 3T MRI scanner with a 12-channel head coil at Aarhus University Hospital, Denmark. The T1-weighted sequence was performed with the following parameters: voxel size 1 x 1 mm, slice thickness 1 mm, matrix size 256 x 256, FoV 256 x 256, repetition time 2000 ms, echo time 3.7 ms, pixel bandwidth 150 Hz/Px, and flip angle of 9°. The diffusion-weighted sequence was acquired using echo-planar imaging (SE-EPI) with a voxel size 2 x 2 mm, slice thickness 2.4 mm, matrix size 96 x 96, FoV 192 x 192, repetition time 5900 ms, echo time 84 ms, b-value 1000 s/mm^2^, 71 diffusion directions (9 *b*0 scans, aquired every 8 volumes), two phase-encoding directions (anterior-posterior and posterior-anterior), pixel bandwith 2004 Hz/Px.

For each participant, probabilistic tractography was applied to the DTI data to produce a map of whole-brain structural connectivity (SC) and thus characterise the strength of physical binding between all brain regions. We used the FDT pipeline in FSL (FMRIB Software Library, Oxford, version 5.0, www.fmrib.ox.ac.uk/fsl/), combined with in-house written scripts, to run the multiple preprocessing steps for SC estimation. These included correction for head movements and eddy currents using the EDDY and TOPUP algorithms with the dual phase-encoding directions, to reconstruct a single set of data with significantly reduced distortions^46^. We fitted a tensor model to each voxel of the brain and visually inspected the coding of the fiber-tracts, followed by estimation of crossing fibers using a Markov-Chain-Monte-Carlo algorithm. Brain parcellation was performed using the Automated Anatomical Labelling (AAL) brain atlas^47^, where the brain is parcellated into 90 cortical and subcortical regions. The AAL atlas is widely used in connectivity studies and can thus facilitate comparison between studies. The estimation of SC was done using probabilistic tractography at the voxel level with a sampling of 5000 streamlines per voxel. For each region, the connectivity to the remaining 89 regions was calculated and normalized by the volume of each region, to control for the area size effect. A threshold of 1% of the maximum number of streamlines from seed to target region was set to remove spurious connections. Since the connectivity probability from region 1 to region 2 is highly correlated with the connectivity probability from region 2 to region 1, we defined the unidirectional connectivity between two regions by averaging these two probabilities. For each participant, a 90 x 90 connectivity matrix was constructed, representing the participant-specific SC network of the brain. Figure 1 illustrates the analysis pipeline.

**Figure 1.**
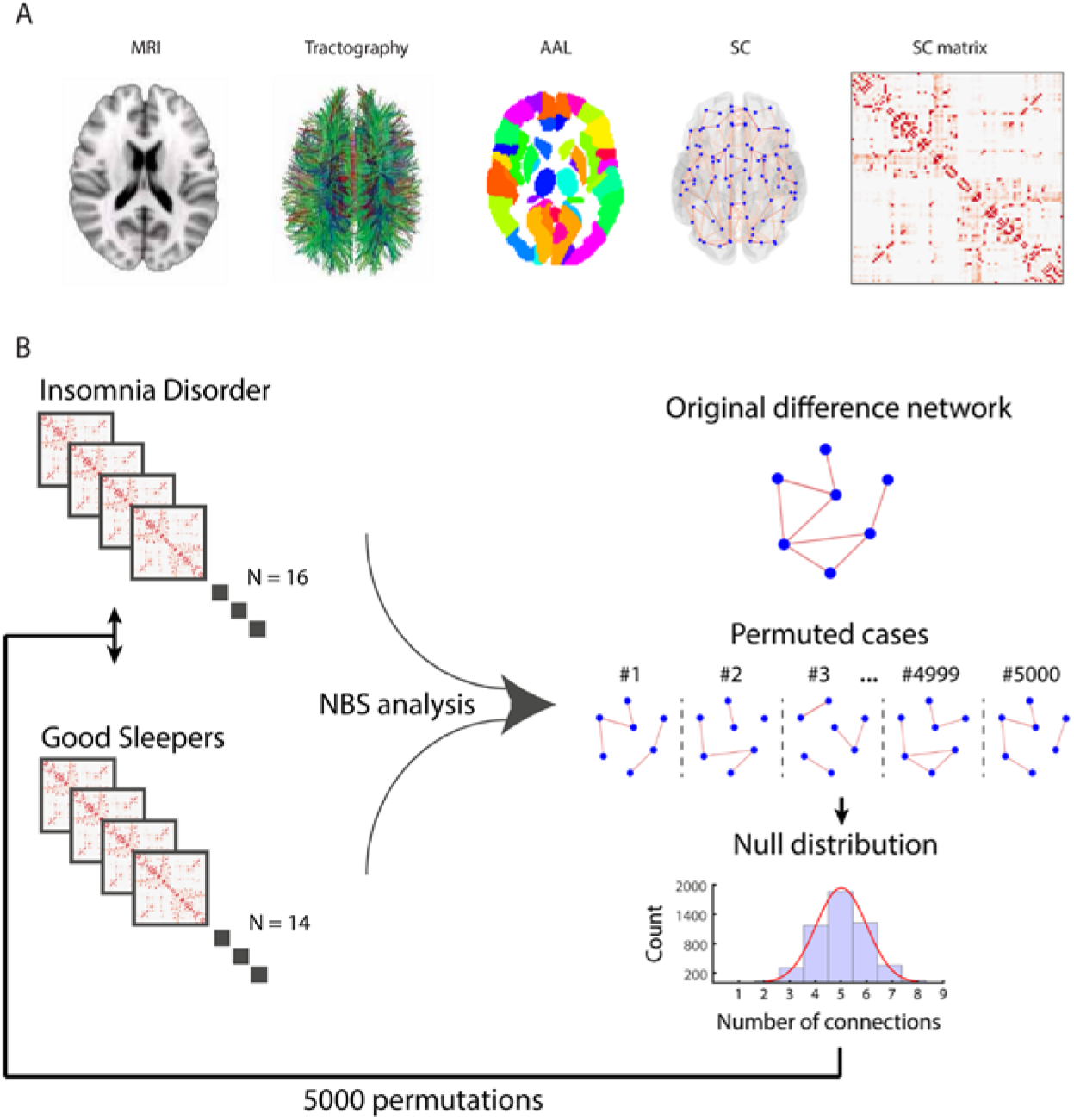
Analysis pipeline. (A) Using standard anatomical MRI and diffusion-tensor imaging, structural connectivity (SC) was determined for each participant by applying probabilistic tractography. The brain regions were defined using the Automated Anatomical Labeling (AAL) template. The SC of each participant can be visualized as a connectivity matrix reflecting the strength of the connectivity between each of the 90 AAL regions. (B) We used Network Based Statistics to compare the connectivity matrices of the two groups. This method creates a null distribution of the largest sub-network by permuting participants across the two groups.

### Group analysis

Between-group statistical comparison was performed using network-based statistics (NBS)^48^. This is a non-parametric method used to identify significant differences between groups of networks, while controlling for multiple comparisons that arise from comparing a large number of connections. This method has been used to assess alterations of brain connectivity in a number of neurological and psychiatric disorders, including schizophrenia^49–51^, ADHD^52–55^, depression^56–60^ and epilepsy^61^. The NBS method controls the family-wise error rate by making the assumption that relevant differences in connectivity between groups are confined to connections that are connected with each other and thus form sub-networks in the larger network. The method is analogous to cluster-based correction strategies used in standard parametric mapping^62^, but rather than clusters in volumetric data it identifies sub-networks in the topological space. NBS is based on the General Linear Model combined with non-parametric permutations of the design matrix. Here, we used a design corresponding to unpaired t-tests between the group diagnosed with Insomnia Disorder and the group of matched controls with no sleep problems. The NBS method relies on a specification of the t-statistic (t-stat) level above which connected sub-networks will be identified, while the permutations control for the family-wise error rate. We tested a range of t-stat thresholds from 2 to 3.4. A high t-stat threshold implies that only the connections with most pronounced differences between groups are included, while lower thresholds allow more subtle differences to form the subnetworks. For all thresholds we regarded sub-networks of corrected p-values as significant if *p* < 0. 05. In all cases, significance was an attribute of the sub-network in question and never individual connections.

### Associations between structural connectivity and clinical scores

We used Pearson’s correlations to assess the relationship between the properties of the identified network and clinical scores of disturbed sleep (PSQI) and insomnia severity (ISI).

## Results

The difference between the groups in PSQI and ISI scores (both p < 0.000) shows that the two groups are clearly distinct with regard to disturbed sleep and insomnia severity. The groups did not differ in terms of age, gender and education (see Table 1). The duration of the sleep problems in the insomnia group ranged from 1 to 20 years with a group mean of 11 years (SD 1.62). According to the polysomnography data, it took around 17 minutes for the participants in the insomnia group to fall asleep. The average total sleep time was 6 hours and 20 minutes, and the mean awake time after sleep onset was 40 minutes. On average, insomnia participants were asleep 85% of the time spend in bed.

### Structural connectivity

Compared to matched controls with normal sleep, the NBS analyses showed that participants with Insomnia Disorder had reduced structural connectivity in a brain network including 34 regions and 39 structural links between them (t-threshold 2.6, p = 0.014). Figure 2 shows a visualization of the network of reduced connectivity in the insomnia group, and Table 2 gives an overview of the difference network ordered by the degree, i.e. the number of connections linking one node to other nodes within the significant network.

**Figure 2.**
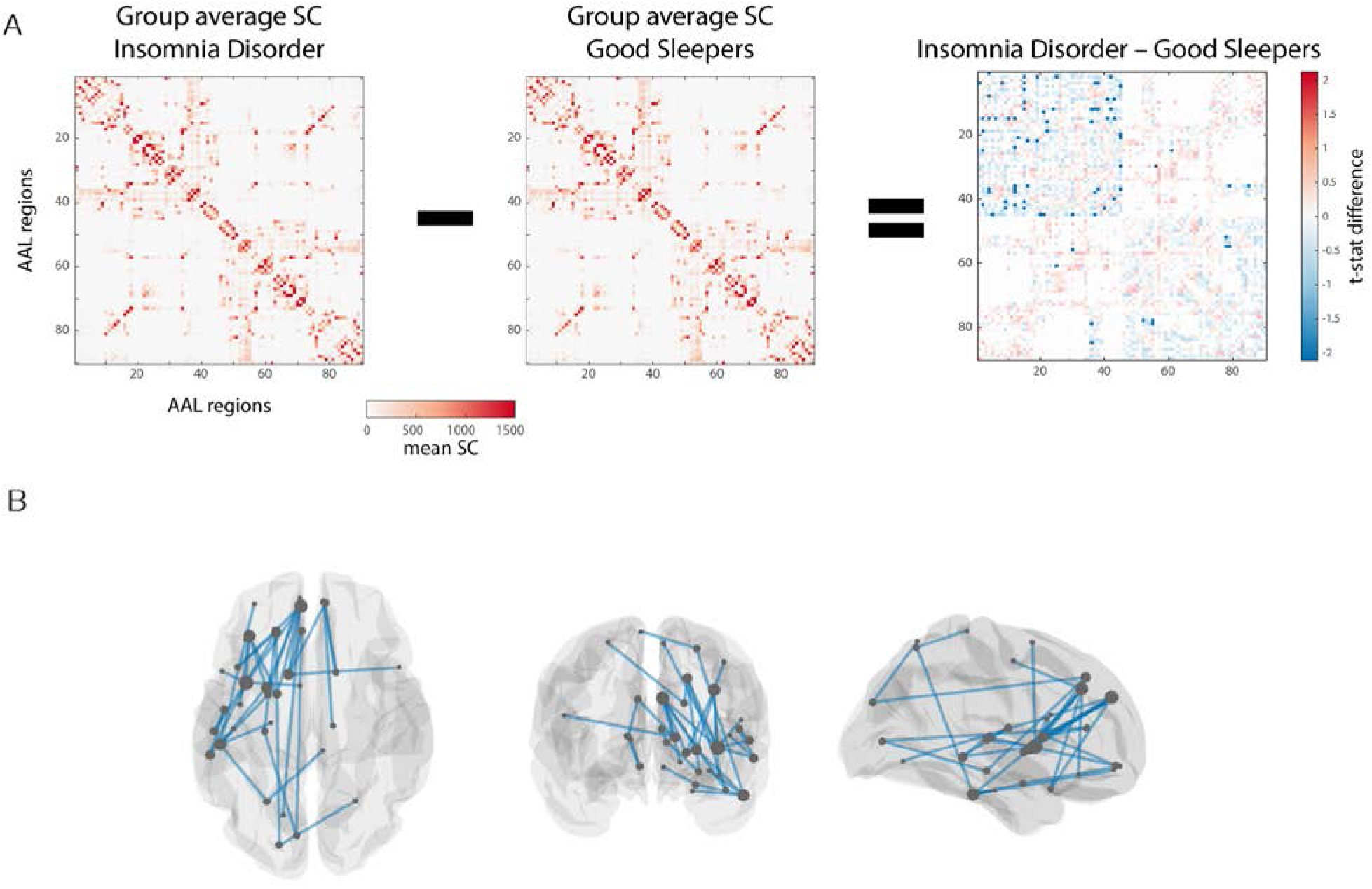
Reduced structural connectivity in Insomnia Disorder. (A) When comparing the connectivity matrices of the participants with Insomnia Disorder to the Good Sleeper Controls, we found a sub-network of significantly reduced connectivity in the insomnia group. (B) A visualization shows the regions and connections involved in the network of reduced connectivity in Insomnia Disorder. Thickness of connection reflects the amplitude of the difference (absolute of t-stat) and node size is scaled to the degree within the significant network.

**Table 2.** Network of reduced structural connectivity in Insomnia Disorder (NBS, t-thresh = 2.6). All connections and their t-value ordered by degree.

| **AAL region 1** | **AAL region 2** | **T-value** |
| --- | --- | --- |
| Insula L | Mid temporal gyrus L | 3.38277 |
| Insula L | Mid frontal gyrus L | 3.31802 |
| Insula L | Sup temporal gyrus L | 3.28373 |
| Insula L | Suppl motor area L | 3.21040 |
| Insula L | Sup frontal gyrus, medial L | 3.03876 |
| Insula L | Sup frontal gyrus L | 2.91902 |
| Insula L | Inf frontal gyrus, operc part L | 2.75371 |
| Sup frontal gyrus, medial L | Putamen L | 3.49883 |
| Sup frontal gyrus, medial L | Ligual gyrus | 2.92450 |
| Sup frontal gyrus, medial L | Temporal pole: sup temp gyrus L | 2.89226 |
| Sup frontal gyrus, medial L | Pallidum L | 2.79188 |
| Sup frontal gyrus, medial L | Caudate L | 2.70861 |
| Mid frontal gyrus L | Putamen L | 3.27051 |
| Mid frontal gyrus L | Precentral gyrus L | 3.12777 |
| Mid frontal gyrus L | Pallidum L | 3.02592 |
| Mid frontal gyrus L | Inf frontal gyrus, orb part L | 2.60041 |
| Inf temporal gyrus L | Putamen L | 3.61412 |
| Inf temporal gyrus L | Sup parietal gyrus L | 3.07406 |
| Inf temporal gyrus L | Amygdala L | 2.86488 |
| Inf temporal gyrus L | Parahippocampal gyrus L | 2.82760 |
| Inf temporal gyrus L | Mid frontal gyrus, orb part L | 2.68540 |
| Sup frontal gyrus L | Sup occipital gyrus L | 3.02766 |
| Sup frontal gyrus L | Temporal pole: sup temp gyrus L | 2.70466 |
| Sup frontal gyrus L | Hippocampus L | 2.66740 |
| Caudate L | Anterior cingulate L | 3.60375 |
| Caudate L | Sup frontal gyrus, medial R | 2.81492 |
| Caudate L | Inf frontal gyrus, operc part R | 2.61736 |
| Putamen L | Pallidum L | 2.83568 |
| Mid temporal gyrus L | Rolandic operculum L | 2.86214 |
| Mid temporal gyrus L | Heschl gyrus L | 2.77894 |
| Caudate R | Sup frontal gyrus, medial R | 2.91264 |
| Caudate R | Sup frontal gyrus, medial orb R | 2.72567 |
| Sup frontal gyrus, med orb R | Thalamus R | 3.67602 |
| Rolandic operculum L | Sup temporal gyrus L | 4.23588 |
| Anterior cingulate L | Calcarine fissure L | 2.74708 |
| Amygdala L | Sup frontal gyrus, medial orb L | 3.05236 |
| Hippocampus L | Calcarine fissure L | 3.09335 |
| Sup occipital gyrus L | Sup parietal gyrus R | 3.04965 |
| Sup parietal gyrus L | Paracentral lobule R | 2.61789 |

AAL = automated anatomical labeling; NBS = network-based statistics; t-thresh = t-threshold; degree = number of connections linking this node to other nodes of the network

The altered network identified with NBS was largely left-lateralized and predominantly involved fronto-subcortical connections. The region of the network with the largest degree was left insula followed by left medial superior frontal gyrus, left middle frontal gyrus, left inferior temporal gyrus, left superior frontal gyrus, left caudate and left putamen (see Table 2). The findings were robust to other NBS settings (see Figure S1 at the end of the manuscript). The NBS analyses revealed no networks of increased connectivity in the insomnia group.

### Correlation with clinical measures

Across all participants, there were significant negative correlations of the mean connection strength of the identified network with the PSQI scores (r=-0.71, p<0.000) and with the ISI scores (r=-0.71, p<0.000) (see Figure 3). There were no significant correlations with the PSQI or ISI scores when looking at correlations within the groups separately.

**Figure 3.**
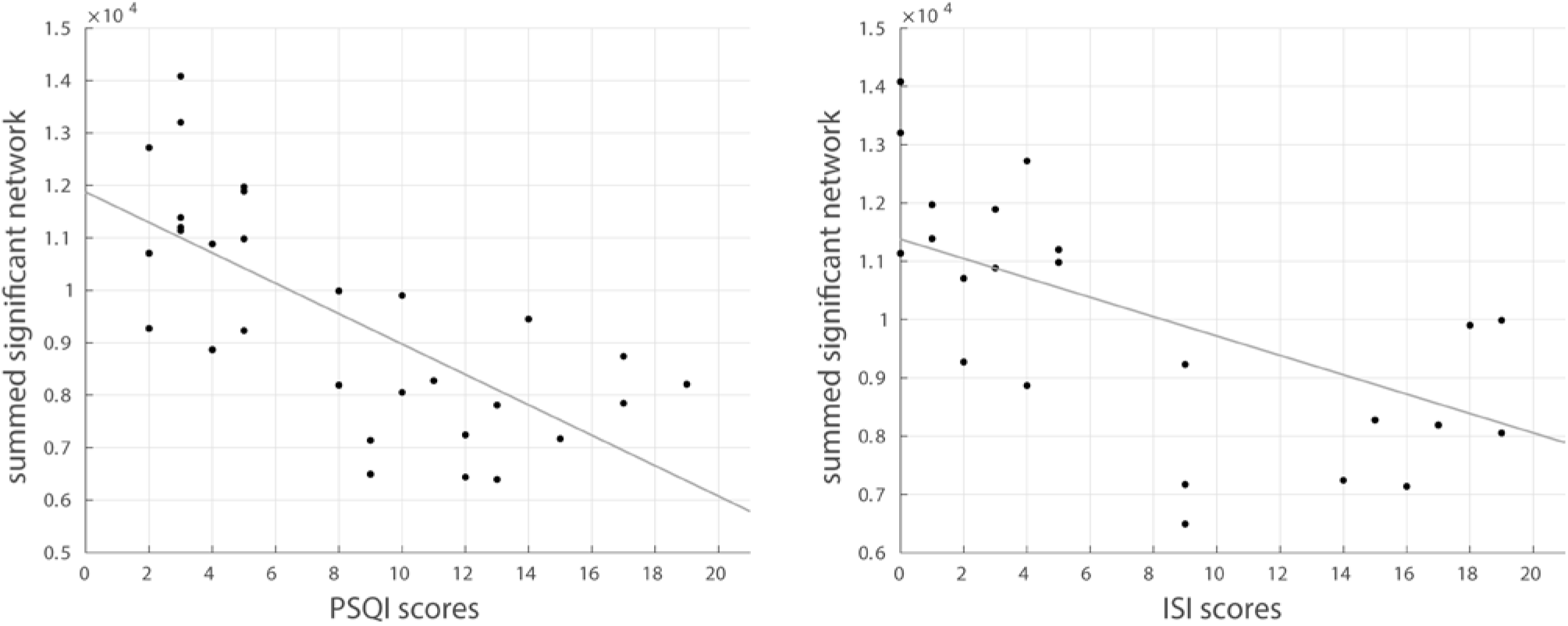
Correlation with clinical measures. Significant correlations were found between mean connection strength of the identified network and clinical measures of sleep quality (PSQI) and insomnia severity (ISI) of all participants. In both scales, higher scores indicate poorer sleep (PSQI) or more severe insomnia (ISI).

## Discussion

We found that Insomnia Disorder is associated with reduced structural connectivity in a network including mainly fronto-subcortical connections in the left hemisphere with particularly impaired connectivity of the left insula. Moreover, we found a clear relationship between mean connection strength of the network and insomnia severity (ISI) and sleep quality (PSQI) across all participants. The findings suggest either that Insomnia Disorder could lead to reduced structural connectivity in fronto-subcortical areas, or alternatively, that people with pre-existing low structural connectivity in these areas have an increased the risk of developing Insomnia Disorder. The mean duration of the insomnia symptoms was 11 years in this sample (range 1 to 20 years), so it would be possible that the disorder could have caused changes in structural connectivity. However, we cannot determine the causality from the present study.

### A key role of the insula

The present findings indicate that reduced connectivity of the left insula plays a key role in insomnia disorder. This is consistent with a number of resting-state functional MRI (fMRI) studies suggesting a role of the insula in insomnia^18,20,23,25,27,63,64^. Reduced functional connectivity between the left insula, amygdala, striatum and thalamus was found in an early resting-state fMRI study comparing persons with Insomnia Disorder to matched controls without sleep complaints^18^. Similarly, Chen and colleagues reported altered insula activation using a resting-state paradigm with simultaneous EEG and fMRI. They found increased co-activation of insula with salience networks in persons with Insomnia Disorder when instructed to fall asleep^20^. The salience networks of this study involve different areas than the connections found in our study, and the relationship between structural and functional connectivity is not simple^65^, but both studies point to insula connectivity as an important factor for understanding the neural circuitry underlying Insomnia Disorder. Furthermore, aberrant insula activity in persons with insomnia was reported in resting-state fMRI studies applying analyses of brain entropy^23^, regional homogeneity^63^ and fractional amplitude of low frequency fluctuations (fALFF)^64^.

Functional neuroimaging studies using positron emission tomography (PET) have also reported altered cerebral glucose metabolism in the insula region^66,67^. For example, Kay and colleagues found that lower sleep efficiency as measured with sleep diaries, was associated to lower relative glucose metabolism in the left insula during wakefulness^67^. Taken together, the results of these previous studies support the findings of the present study suggesting disrupted insula connectivity in Insomnia Disorder.

Structural neuroimaging studies have also pointed to insular involvement in insomnia complaints. A recent DTI study of healthy adults with insomnia symptoms reported a negative correlation between nodal efficiency in the left insula and insomnia severity^39^. Nodal efficiency is a graph theoretical term reflecting how well a brain region connects with other regions of the network. As such, these results seem in line with our findings, even though the study provides no information on the specific nature of the connections. In addition, a voxel-based morphometry study reported an inverse association of early morning awakening with grey matter density in the area bordering the left insula and orbitofrontal cortex^32^.

The insula is thought to play an essential role for human awareness, and it is involved in a number of functions, such as interoception, emotional salience, time perception and decision-making^68,69^. Studies suggest that insula is important for integration of information on our feeling state^70^. It is involved in visceral and somatic sensory processing and the integration of these processes into conscious emotional experience^69,71^. Insular activity is particularly linked to interoceptive awareness^72^, and recent studies showed alterations in EEG-markers of interoception and somatic awareness in Insomnia Disorder^73,74^. The reduced insula connectivity observed in this study may be key to altered interoception and somatic awareness in Insomnia Disorder.

Insula function is not limited to interoception and affective processes. It is also a key region in the detection and processing of salient stimuli across multiple cognitive and sensory domains^71^. The insula is uniquely positioned deep within the lateral sulcus of the brain, widely connected to frontal, temporal, parietal and subcortical areas, particularly amygdala and thalamus^75^. This position enables a key function in the so-called salience network involved in the orientation towards both external and internal stimuli and the generation of appropriate responses to these^76^. These responses also include the modulation of autonomic reactivity to salient stimuli^71,76^, and as such disturbed insula connectivity may be a primary target for studies on brain mechanisms underlying hyper-arousal, the enigmatic key feature of Insomnia Disorder. In summary, the reduced connectivity of the insula suggests that insomnia is associated with impairments in interoception, emotional processing and the generation of appropriate internal and external responses to salient events.

The insular cortex is very closely apposed to the claustrum, a thin subcortical sheet of neurons. Given their proximity, we cannot exclude a possible involvement of claustrum connectivity where we interpret connections as having the insular cortex as origin or destination. Interestingly, a recent genome-wide association study, combined with cell type-specific gene-set analyses, showed significant enrichment of insomnia risk genes in pyramidal neurons of the claustrum^40^.

In a meta-analysis, insula grey matter reductions have been found as a shared feature of psychopathology, including depression, anxiety and schizophrenia^77^. This is consistent with the present results since sleep disturbances are highly prevalent in most psychiatric disorders^10,78^. Neuroimaging studies investigating these should take care to control for the impact of insomnia. The exclusion criteria of the present study ensured that participants did not suffer from psychiatric disorders. Still, the question remains if insula dysfunction is in any way specific to insomnia or may be a more general vulnerability factor that is involved in a number of diseases. A recent large scale study reported a role of the insula in the functional connectivity structures mediating the association between depressive symptoms and sleep quality^79^, and future studies should aim at disentangling the role of insula in insomnia and psychiatric disorders.

We found that participants with Insomnia Disorder showed a network of reduced connectivity with the left insula being connected to seven other regions within this network (primarily frontal regions, see supplementary table 1). It would be relevant to know if these connections are afferent, efferent or bi-directional connections, but it is not possible to determine the nature of the connections from our data. In general, we still have a limited understanding of the structural connectivity of the human insula, even though recent progress has been made^80^. Therefore, we need future research to assess whether Insomnia Disorder is related to the output of the insula not reaching downstream areas sufficiently or if the input to the insula is impeded and therefore insula activity is not modulated optimally.

### Reduced fronto-subcortical connectivity in Insomnia Disorder

Another key feature of the present findings is the reduced fronto-subcortical connectivity. Fronto-subcortical network dysfunction has been identified in several neurological and psychiatric disorders^81^, and it is also consistent with previous studies on Insomnia Disorder. Studies of grey matter changes in insomnia reported reductions in frontal areas, including the left orbitofrontal cortex^31^ and the dorsolateral and medial prefrontal cortex^82^. Previous studies looking at white matter integrity also found reduced fractional anisotrophy in the anterior internal capsule in persons with Insomnia Disorder compared to good sleeper controls, indicating disturbed fronto-subcortical connectivity^35,36^. Our results are also in line with a recent DTI study that found reduced fronto-subcortical connectivity in Insomnia Disorder^37^. However, this study also reported other networks of reduced structural connectivity and one network of increased connectivity. The reasons for the divergence of the results may be found in the data quality of the study, the use of deterministic tractography and unclear NBS settings.

The reduced connectivity in the present study includes connections between frontal areas and classical limbic areas, such as amygdala, hippocampus and thalamus, and alterations in these regions have previously been reported in persons with Insomnia Disorder^18,33,34^. However, what is more pronounced is the reduced connectivity of frontal regions, including the orbitofrontal and medial frontal cortex with basal ganglia structures, such as the putamen, caudate and pallidum. The basal ganglia are a group of subcortical nuclei involved in a range of functions, including motor control, reward-based learning and working memory^83^. A recent genome-wide association study, combined with tissue-specific gene-set analyses, showed strong enrichment of insomnia risk genes across the basal ganglia^40^, and altered activity in basal ganglia structures has previously been associated with Insomnia Disorder. A recent study reported that structural changes of the putamen was related to higher arousal indices in participants with persistent insomnia^34^. Likewise, differences in the local topology of the putamen were found to be related to increased insomnia scores in a recent study using resting-state fMRI in healthy adults with insomnia symptoms compared to controls with no insomnia complaints^24^. In a task-based fMRI study Stoffers and colleagues found reduced recruitment of the head of the left caudate in persons with Insomnia Disorder during an executive task and argued that the caudate was an essential structure for the abnormalities related to insomnia^84^. This suggestion is supported by a seed-based resting-state fMRI study by Huang and colleagues, which had amygdala as primary region of interest. They reported reduced connectivity between left amygdala and bilateral caudate as well as between the right amygdala and the left pallidum^18^. In summary, these studies are consistent with our findings that point to an essential role of the basal ganglia in Insomnia Disorder.

A recent meta-analysis of neuroimaging studies on post-traumatic stress disorder (PTSD) found that persons with PTSD differed from trauma-exposed persons without PTSD mainly in altered activity in basal ganglia regions^85^. Interestingly, pre-existing insomnia symptoms likewise increase the odds of developing PTSD after trauma exposure^86^. It may be that basal ganglia dysfunction is involved when stress responses turn into a clinical syndrome, such as PTSD. Our results suggest that reduced basal ganglia connectivity with frontal regions is an important component of the pathophysiology of insomnia and the risk it imposes on the development of other mental disorders.

### Left hemisphere lateralization

A prominent feature of the present findings is the high degree of left hemisphere lateralization. The left hemisphere has not previously been emphasized in relation to insomnia as most studies find bilateral alterations of brain activity. The lateralization is not explained by handedness, as we ran the NBS analysis with handedness as covariate and found the same results. Interestingly, mostly left hemisphere alterations in insomnia disorder or in association with insomnia symptoms have been reported previously, including reduced volume of the left orbitofrontal cortex^31^ and of the area bordering the left insula and orbitofrontal cortex^32^ as well as reduced activation in the head of the left caudate during executive tasks^84^. Reduced nodal efficiency of the left insula and putamen has also been associated with Insomnia Disorder^37^. These previous findings all show reductions in left hemisphere structures that are part of the network of reduced connectivity found in the present study.

Furthermore, altered functional connectivity of the left insula seems to be one of the most consistent findings of resting-state fMRI studies^18,20,63,64^. Based on asymmetries in the peripheral autonomic nervous system, a neurobiological model has proposed that the left insula is primarily activated by parasympathetic activity and the right insula is activated predominantly in relation to sympathetic input^87^. According to this model, reduced structural connectivity of the left insula, as seen in our results, would be expected to impair the neural processing of parasympathetic input and as such interfere with sleep, since non-REM sleep is characterized by a decrease in sympathetic activity and increased parasympathetic activity^88,89^.

In a different line of work, high-density EEG research on slow waves during sleep has shown a higher involvement of the left hemisphere in the origin and propagation of slow waves during sleep^90^. Particularly, slow waves often originate in the left insula and involve the middle, medial and inferior frontal gyri significantly more in the left hemisphere than in the right. These areas are exactly the key regions of reduced connectivity identified in our study. An extensive meta-analysis shows that persons with insomnia have significantly reduced slow wave sleep compared to controls with no sleep problems^91^, and the network of reduced connectivity identified in this study may be one of the mechanisms behind the reduction in slow wave sleep in insomnia. Furthermore, a recent study shows that the sleep EEG of Insomnia Disorder is characterized most by a difficulty to transition from N2 sleep to the deeper sleep stage N3 characterized by slow waves^92^. Concertedly, these previous findings and the current results make it tempting to suggest involvement of the reduced connectivity of the left insula and frontal regions in persons with Insomnia Disorder in their difficulty to enter slow-wave sleep.

In the interpretation of the present results, a number of limitations should be considered. First, this is a cross-sectional study, and we can draw no causal conclusions from the results regarding whether the observed differences in structural brain connectivity are the cause or the result of insomnia disorder. Future studies should use longitudinal designs to clarify the causal direction of structural alterations in Insomnia Disorder. Second, the present study included a relatively small sample size. The two groups, however, are well-matched and at the same time clearly distinct with regard to insomnia measures. In addition, the insomnia group is very well-defined, as all these participants group underwent polysomnography to rule out common comorbid sleep disorders such as sleep-disordered breathing and sleep-related movement disorders^93,94^. Finally, we could not assess subdivisions of the relevant network regions, such as anterior or posterior insula, since the AAL atlas used for brain parcelation includes only relatively large brain regions. We chose this atlas because it is commonly used and thereby facilitates comparison between studies, but a different brain parcellation strategy might allow for a more nuanced view of the brain regions.

In conclusion, our results show that Insomnia Disorder is related to significantly reduced structural connectivity in a network involving left insula as well as fronto-subcortical connections. This study adds to the existing knowledge by taking a data-driven whole-brain perspective on brain connectivity. The findings demonstrate changes in brain connectivity at the structural level that form the anatomical ‘backbone’ of functional connectivity in areas related to interoception, emotional processing, stress responses and the initiation of travelling slow waves. These findings can improve the understanding of the neurobiological foundations of insomnia and may assist the development of efficient treatment strategies.

## Acknowledgements

Thanks to Tim Van Hartevelt, Dora Zeidler, Marit Otto and Kristina Bacher Svendsen for assistance with this project. Center for Music in the Brain is funded by the Danish National Research Foundation (DNRF117).

## Abbreviations

AAL: Automated Anatomical Labelling Brain Atlas
DTI: Diffusion Tensor Imaging
fMRI: Functional Magnetic Resonance Imaging
ISI: Insomnia Severity Index
NBS: Network Based Statistics
PSQI: Pittsburgh Sleep Quality Index
PTSD: Post-Traumatic Stress Disorder
REM: Rapid eye movement sleep
SC: Structural connectivity
SD: Standard deviation

## Supplementary figure S1

**Figure S1 –.**
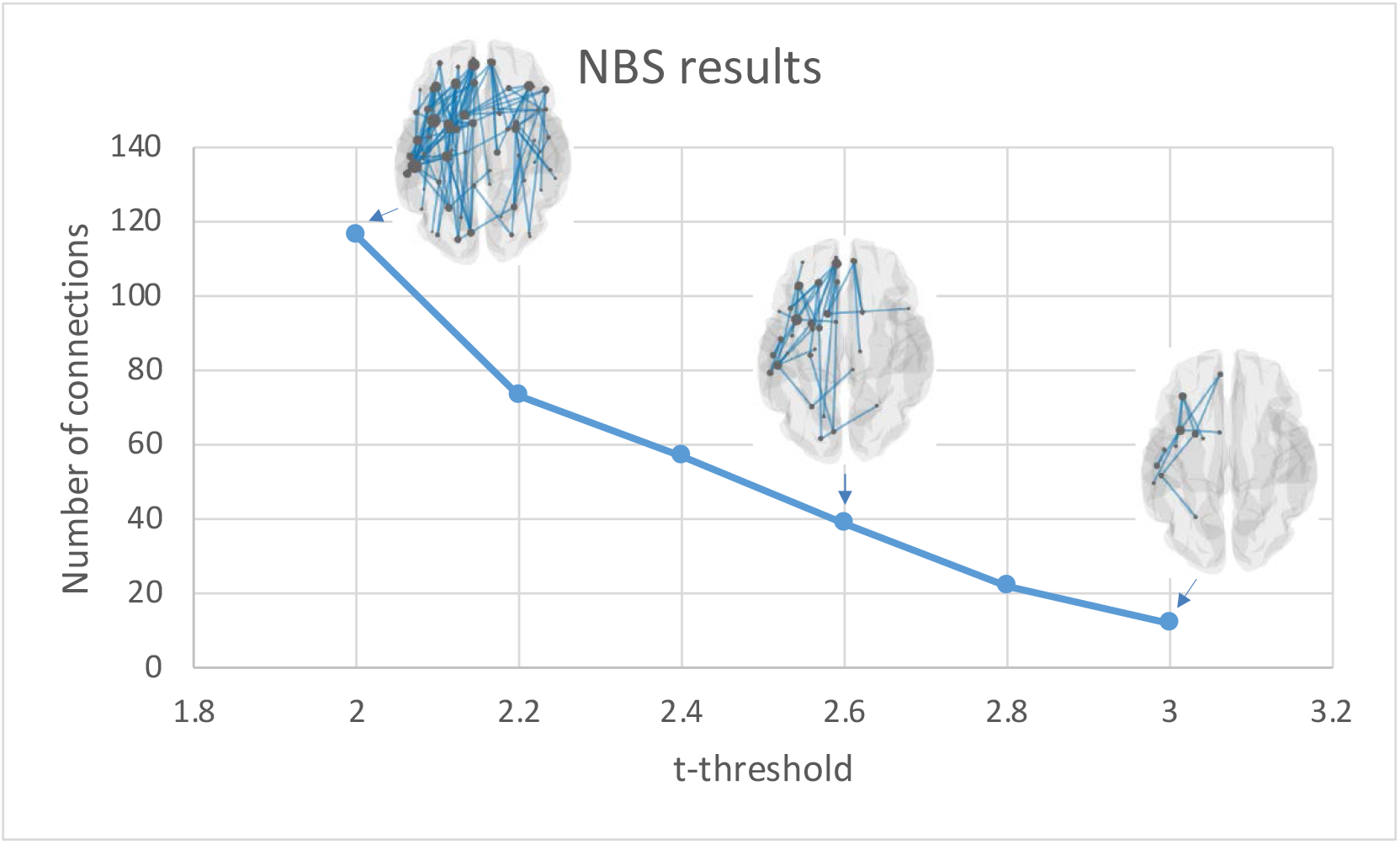
NBS settings. This figure shows the significant networks identified at a range of t-stat thresholds with the Network-based Statistics (NBS). We tested the NBS analyses at a range of thresholds (2, 2.2, 2.4, 2.6, 2.8, 3, 3.2 and 3.4), and similar significant networks were identified in the threshold range 2 to 3. Lower t-thresholds are associated with a larger number of connections whereas larger t-thresholds include only the connections with the most pronounced difference between the two groups. There were no significant results for the t-thresholds 3.2 and 3.4. At the largest significant t-threshold of 3, the network included 12 nodes with 12 connections between them. Ordered by degree, these regions were the insula, middle frontal gyrus, putamen, superior frontal gyrus (medial part), superior temporal gyrus, inferior temporal gyrus, pallidum, rolandic operculum, supplementary motor area, precentral gyrus, middle temporal gyrus and the superior parietal gyrus all in the left hemisphere. The threshold of 2.6 was chosen as the optimal balance between detail and overview.

